# First attempt to validate multiplex PCR with sex markers SSM4 and ALLWSex2 in long-term stored blood samples of eastern North American shortnose sturgeon (*Acipenser brevirostrum*)

**DOI:** 10.1101/2024.10.31.621305

**Authors:** Hajar Sadat Tabatabaei Pozveh, Salar Dorafshan, Tillmann J. Benfey, Jason A. Addison, Matthew K. Litvak

## Abstract

Sex-specific information is crucial for sturgeon culture, conservation, and fisheries management. However, identifying the sex of sturgeon is difficult, especially for immature individuals. Two recent studies identified two female-specific loci (AllWSex2 and SSM4) that are conserved among many Acipenserid species, but they have not been validated for all species within this family. The objectives of this study were to 1) determine whether SSM4 can be used to sex shortnose sturgeon; 2) develop and test a multiplex PCR technique using both ALLWSex2 and SSM4 for sexing shortnose sturgeon; 3) determine if long-term storage of blood samples can be used to sex shortnose sturgeon; and 4) test the effect of storage temperature on DNA degradation. DNA was extracted from frozen RBC samples from 36 fish which had previously been sexed. A multiplex PCR was set up using three pairs of primers: AllWSex2 and SSM4, as female-specific loci, and mtDNA as an internal control which were all run on a 2 % agarose gel. AllWSex2 and SSM4 allowed for perfect discrimination of sex. While there was DNA degradation, as a result of long-term storage and temperature, the signal was still strong enough after 8 years of cold storage to delineate sex. This suggests that researchers now have the ability to reexamine archived/frozen samples to determine sex of their fish.

## 1. Introduction

*Acipenserids*, which have existed for over 250 million years [1], are among the most threatened vertebrate group listed by the International Union for Conservation of Nature [2]. All 27 living species of *Acipenserids* (sturgeon and paddlefish) are currently listed; of these, 17 are in decline, 4 are stable, 3 are of unknown population status and 2 are increasing (IUCN, 2023). Globally, sturgeons are threatened by the combined pressures of overfishing, habitat degradation, river fragmentation, and poaching [3,4]. Sturgeons are the only fish that produce caviar, one of the most highly valued animal products worldwide [3,5]. Wild sturgeons are long-lived, late to mature, and have long reproductive cycles [6–8], making it difficult to recover naturally from anthropogenic threats. As global populations of wild sturgeon continue to decline, there is an urgent need to understand their population sex ratio and spawning biomass to inform conservation policy. Early identification of sex is also important for aquaculture as the value of caviar far exceeds that of the meat, and it is therefore more profitable to only grow females.

Sex determination in sturgeon is made difficult by the lack of easily identifiable sexual characteristics in juveniles [9]. However, ultrasound, endoscopy, biopsy, and steroid hormone analysis have all been used, with varying success, for sexing older sturgeons [10], and recently geometric morphometrics were used to discriminate between sexes in mature adult shortnose sturgeon with a high degree of accuracy [11]. There is a predictable relationship between steroid hormone levels and sex in numerous species of sturgeon, including Russian sturgeon (*Acipenser gueldenstaedtii*) [12], stellate sturgeon (*A. stellatus*) [12,13], Atlantic sturgeon (*A. oxyrinchus*) [14], lake sturgeon (*A. fulvescens*) [15], and white sturgeon (*A. transmontanus*) [16]. Matcshe and Gibbons [17] assessed a variety of hematological, plasma chemistry, and sex hormone approaches in a dam-impeded spawning run of shortnose sturgeon (A. *brevirostrum*). They found a difference in levels of sex hormones among mature adults. While these approaches help to delineate sex, none is definitive, and none works on young fish.

More recently, the discovery of genetic markers ALLWSex2 [18] and SSM4 [19] has allowed the determination of sex with a high degree of accuracy for many species of sturgeon, with AllWSex2 used to sex North American species, including shortnose sturgeon [20], lake sturgeon [21–23], gulf sturgeon (*A. oxyrhinchus desotoi*) [20], and Atlantic sturgeon [18,20], and SSM4 used to sex several European sturgeons and Atlantic sturgeon [19], but not shortnose sturgeon. These are great advances and important to our understanding of sex ratios of sturgeon in the wild and under aquaculture production.

Fin clips were used as the source of DNA for the ALLWSex2 studies whereas blood samples were used for the SSM4 study. Collecting fin clips is standard practice in fish ecology, where they have also been used to identify species and as a source of material for stable isotope analysis. However, there are some issues associated with using fin clips for genetic studies. For instance, surgical tools can become contaminated if not cleaned and sterilized properly between samples and there is a potential for contamination through gloves or from the water if multiple fish are held in the same tank. This is not a concern with blood collection as the needles and syringe barrels are clean and sterile. Syringes can be held on ice and then taken back to the lab where nucleated red blood cells can be obtained by centrifugation prior to freezing.

Alternatively, blood can be dropped onto an FTA card to stabilize DNA prior to processing [24,25].

In North America, shortnose sturgeon have been listed as endangered within the U.S. since 1967 [26] and a Species of Special Concern in Canada since 1980 [27]. The shortnose sturgeon’s northernmost population occurs in the Wolastoq River (aka Saint John River, NB), its only known location in Canada. Like other sturgeons, shortnose sturgeon populations are threatened by river pollution, fragmentation due to damming, angling practices, and bycatch from other commercial and recreational fisheries [27–32]. Current assessments of population size, sex ratios, frequency of spawning, and spawning biomass are key to the effective management of this species, especially when creating policies for commercial and recreational fishing [27]. Unfortunately, there has been limited research done to assess the total river population, reproductive spawning biomass, or sex ratios of shortnose sturgeon in the Wolastoq River since Dadswell [28]. Clearly, there is a need for accurate and non-invasive techniques to determine sex ratio of this population and aquaculture development in North America.

The goals of this study were to 1) determine whether SSM4 can be used to determine the sex of shortnose sturgeon; 2) develop and test a multiplex PCR technique using both ALLWSex2 and SSM4 for sexing shortnose sturgeon; 3) determine if long-term storage of blood samples can be used to sex shortnose sturgeon; and 4) test the effect of storage temperature on DNA degradation.

## 2. Methods

### 2.1 Sampling and storage

Shortnose sturgeon samples were obtained from an archive in Litvak’s lab at Mount Allison University (Sackville, NB, Canada). All fish from this archive were anesthetized with 150 mg/L MS222 buffered with CaCO_3_. An 18-gauge Luer lock 5-10 cc heparinized syringe was used to withdraw blood while fish were in a supine position on a v-trough table. The caudal vein was accessed by inserting the needle posterior to the anal fin at a 45° angle. Blood was withdrawn using the syringe after a flash of blood was observed in the Luer lock. Syringes were then stored on ice prior to centrifugation. Blood samples were then divided into 2 mL vials and spun in a mini centrifuge (Fisher Scientific) for a minimum of 2 minutes at 2000 g to separate red blood cells (RBCs) from the blood plasma. The RBCs were held on ice, then either flash frozen and held in a liquid nitrogen dry shipper or in a mini-fridge freezer in the field until they were transferred to a -20°C or -80°C freezer in the lab.

### 2.2 Sex confirmation

Blood samples were taken from spermiating males during the spring spawning runs in 2016 and 2022. Litvak’s lab also has ultrasound images of all fish sampled within their tissue archive, and we selected blood samples from females in this collection that clearly showed eggs in ultrasound images (Figure 1). The ultrasound image library was constructed from images taken with either a Sonosite MTurbo or EdgeII ultrasound equipped with a HPLF50 linear probe (5-15Mhz). All fish were anesthetized with 150 mg/L MS222 buffered with CaCO_3_ and then intubated with running water while on a V-trough or held in water in a tank during ultrasonography.

**Figure 1.**
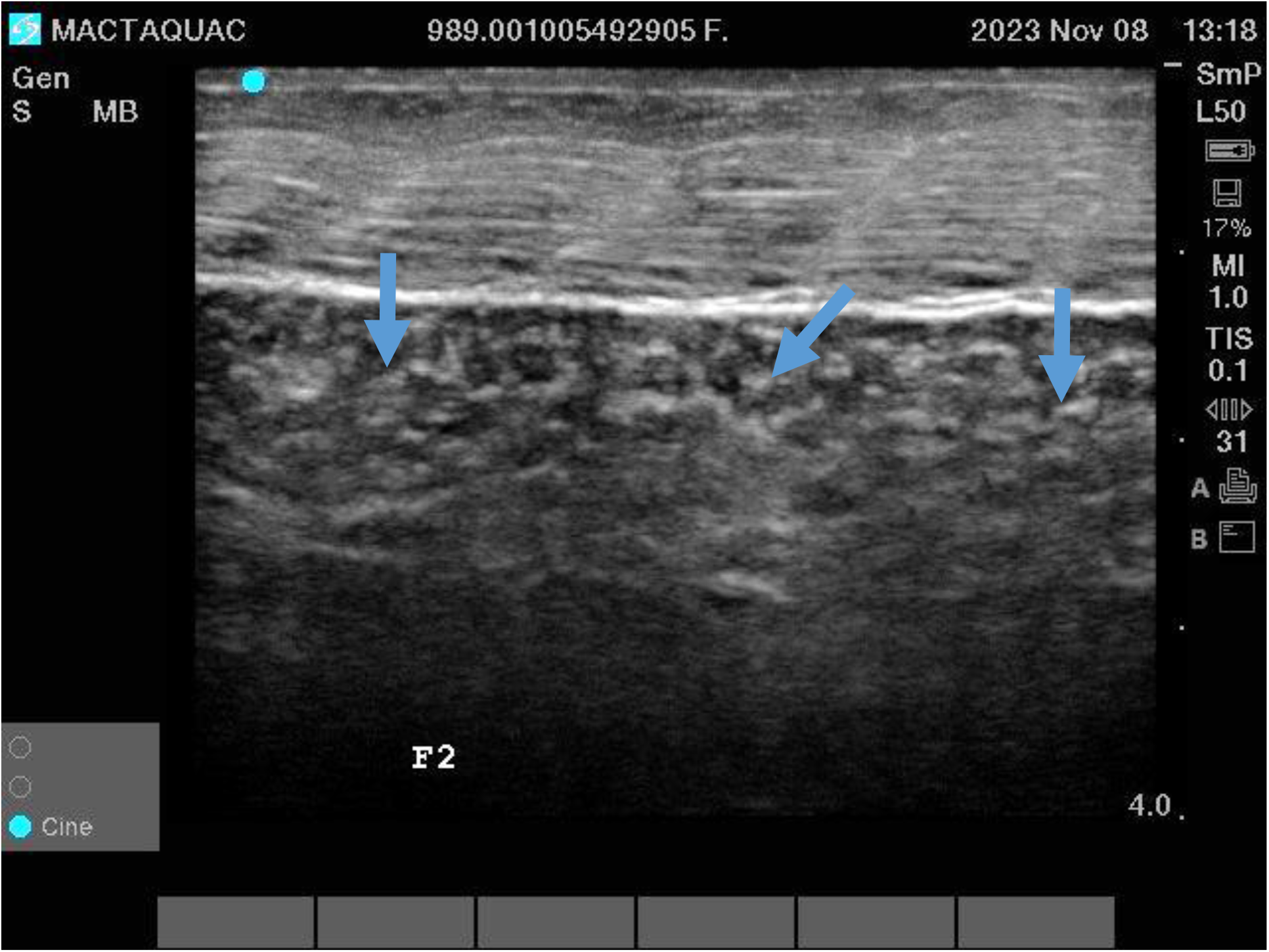
Example of a left lateral frontal ultrasound image of an adult female (FL 80 cm; 3.7 kg) clearly showing the presence of eggs (blue arrows). Image taken with a Sonosite Edge II Ultrasound using a high frequency linear probe (HLF50). While the ultrasound image was taken in November 2023, the blood sample and measurement from this fish were collected in 2016.

### 2.3 DNA yield and purity

We used the DNeasy Blood and Tissue Kit (Qiagen, Hilden, Germany) method for extraction of DNA from the RBC samples. The concentration of DNA was determined using a NanoVueTM UV/Visible Spectrophotometer (GE Healthcare Bio-Sciences Corp) in accordance with the manufacturer’s instructions. Each sample was measured twice, and any inconsistent measurements were repeated a third time. The presence of DNA was verified by using a 1% agarose gel stained with Safe stain. Specifically, 5 μL of each DNA sample was mixed with 3 μL of 10x loading buffer (50% (v/v) glycerin, 0.1 M EDTA pH 8.0, 0.25% (w/v) bromophenol blue, and 0.25% (w/v) xylene cyanol) and subjected to electrophoresis. The DNA appeared as a smeared band under UV light, and its integrity was recorded.

### 2.4 Application of AllWSex2 and SSM4 primers for sex discrimination

The PCR process was initially carried out separately using two pairs of primers, AllWSex2 and SSM4, which are specific for sex determination in shortnose sturgeon. In the subsequent step, a mitochondrial primer pair was optimized for use as an internal control. An internal positive control can be employed to detect the presence of PCR inhibitors. This is achieved through a multiplex reaction in which the target sequence is amplified with one primer set, while a control sequence (i.e., the internal positive control) is amplified with a different primer set (Kuhl et al. 2021; Ruan et al. 2021).

Multiplex PCR was conducted to evaluate the genotypes of AllWSex2 and SSM4 according to the methods outlined by Kuhl et al. (2021)[18] and Ruan et al. (2021)[19], respectively. Primer pair sequences for AllW2 (F: 5′TGATCAACCTCTTCAGCAATGTC3′, R: 5′TGAGAGCCACTGTACTAACACA3′) were utilized to amplify a 120 bp fragment, while primer pair sequences for SSM4 (F: 5′TCGGTATCTTAAACTGAACCAA3′, R: 5′AGATGGAGAATTCATTGCCTA3′) were used to amplify a 400 bp fragment. A 300 bp mtDNA fragment served as an internal control (F:5′CCCTGATCCTAATGTTTTCGGTTGG3′, R:5′AGATCACGTAGGACTTTAATCGTT3′).

The PCR reaction mixture contained 5 pM of each primer and 25 μL of Qiagen Mastermix in a final volume of 50 μL, supplemented with 15 mM MgCl_2_ (Qiagen, Germany). PCR cycling conditions consisted of an initial denaturation step at 95 °C for 10 min, followed by 30 cycles of denaturation at 95 °C for 60 s, annealing at 56 °C for 60 s, and extension at 72 °C for 60 s, with a final extension step at 72 °C. Genotypes of the PCR products were determined by electrophoresis on a 2% agarose gel. We compared the sex identifications derived from the AllWSex2 and SSM4 loci with phenotypic characteristics for each fish, i.e., spermiation for males and ovarian ultrasonography for females (Figure 1).

### 2.5 Experimental design

To remove any potential bias, archived RBC samples from Mount Allison were given a new ID number prior to shipping on dry ice to the University of New Brunswick (Fredericton, NB, Canada) for analysis. Thirty-two samples were analyzed: 24 males and 8 females held for either 2 or 8 years and at either -20°C or -80°C (Table 1).

**Table 1:**
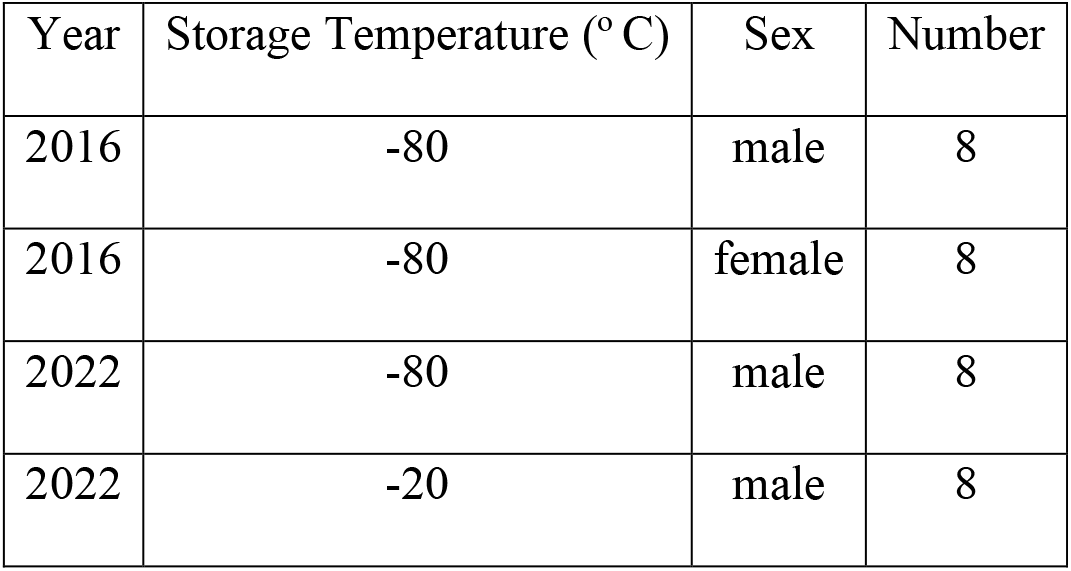
Sample year, RBC storage temperature, sex, and number of fish used for analyses.

We compared the sex assignment, DNA concentration, and the A260/A280 and A260/A230 DNA ratios. The RBCs from the eight females and eight males caught in 2016 and stored at -80°C were used to test the efficacy of the Multiplex PCR approach to determine sex of shortnose sturgeon. We also used these fish to determine the effect of sex on DNA concentration and the two DNA ratios. DNA is ideally extracted immediately after sampling or stored at sub-zero temperatures and extracted within 24 hours [33]. Unfortunately, this is not always possible in the field. We therefore tested the effect of storage temperature and time on DNA degradation with this dataset. Simple t-tests were used to test the differences between these response variables.

## 3. Results

Initially, ALLW primer bands with a size of 120 bp were observed in female individuals (Figure 1, numbers a-c). Subsequently, the SSM4 primer results were analyzed, and female individuals with a band length of 400 bp were observed. The mitochondrial primer used exhibited a band length of 300 bp in all individuals. Subsequently, all three primers were included in a single reaction.

All fish showed the 300 bp mtDNA band, confirming sample integrity, i.e., DNA was present in the samples. The female-specific AllWSex2 (120 bp) and SSM4 (400 bp) alleles were present in all female samples and absent in all males (Figure 2). All phenotypic and genotypic sex identifications agreed, demonstrating 100% accuracy of our multiplex PCR approach. The AllWsex2 primer used in this study previously generated fragments of approximately 100 bp in Kuhl et al.’s ([18]) research. However, in our study, it amplified fragments of around 120 bp.

**Figure 2.**
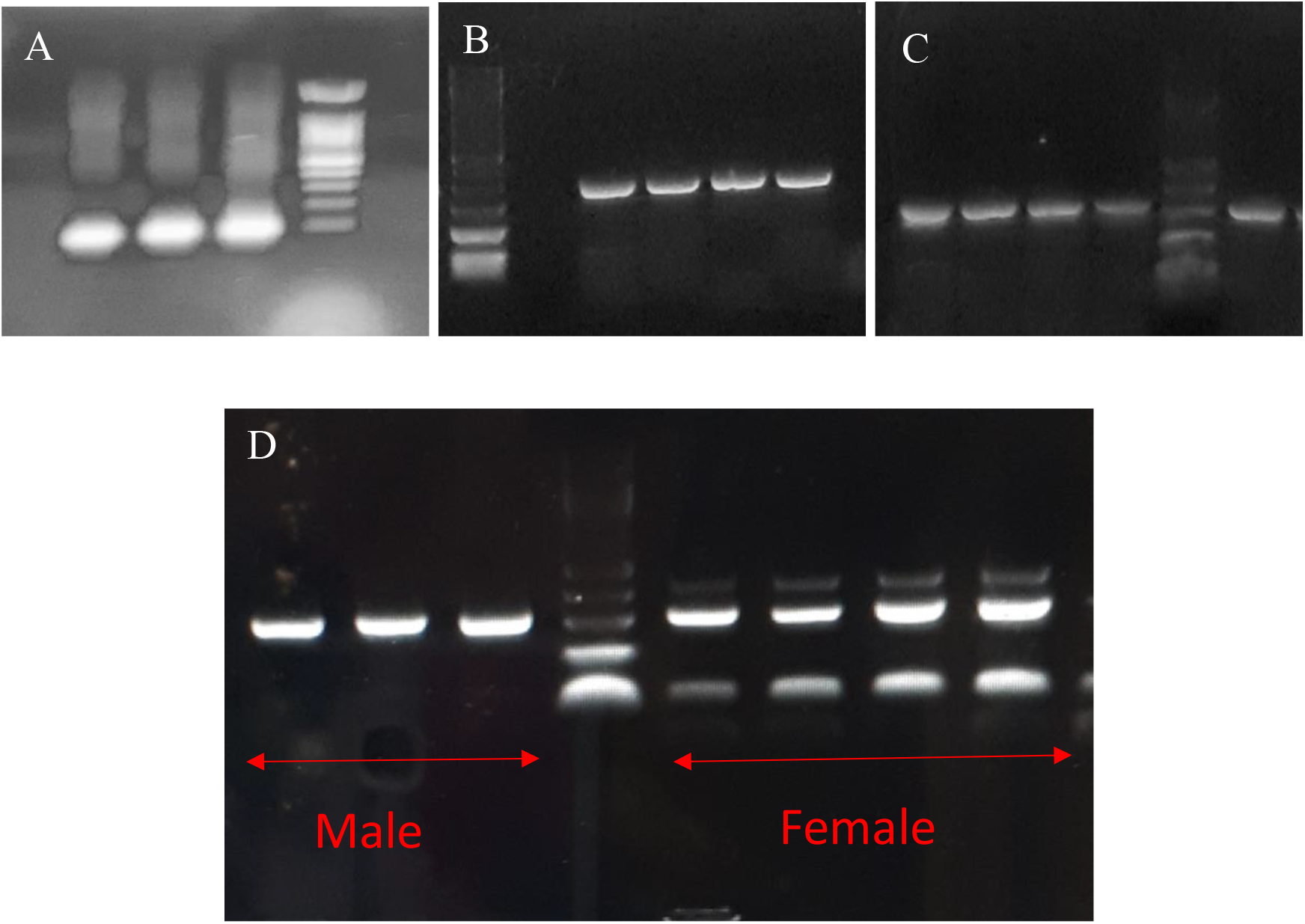
Female-specific markers in shortnose sturgeon using simplex (A, B and C) and multiplex PCR (D). A: Validation of AllWsex2 (100 bp); it was only amplified in females, B: SSM4 (400 bp); it was only amplified in females, C: The internal control (mtDNA; 300bp) was amplified in both males and females, D: Validation of female-specific markers and mtDNA in shortnose sturgeon using multiplex PCR. Ladder is in 100 bp increments.

There was no effect of sex on DNA concentration and A260/A280 or A260/A230 DNA ratios (all three t-tests, df=14, p>0.80; Figure 3) for RBCs held at -80°C since 2016. Since there was no effect of sex on DNA variables, we combined DNA data from males and females held at -80°C to examine the effect of storage time on DNA degradation. Not unexpectedly, there was a significant effect of length of storage time on DNA concentration and ratios (all three t-tests, df=22, p<<0.001; Figure 4). While we did see significant DNA degradation over the 8 years in storage compared to samples held for 2 years, the multiplex PCR technique was still able to delineate sex perfectly. While we were able to successfully discriminate between sexes using RBCs stored at both -20°C and -80°C, there was a significant reduction in DNA concentration and ratios (all three t-tests, df=14, p<0.002; Figure 5).

**Figure 3.**
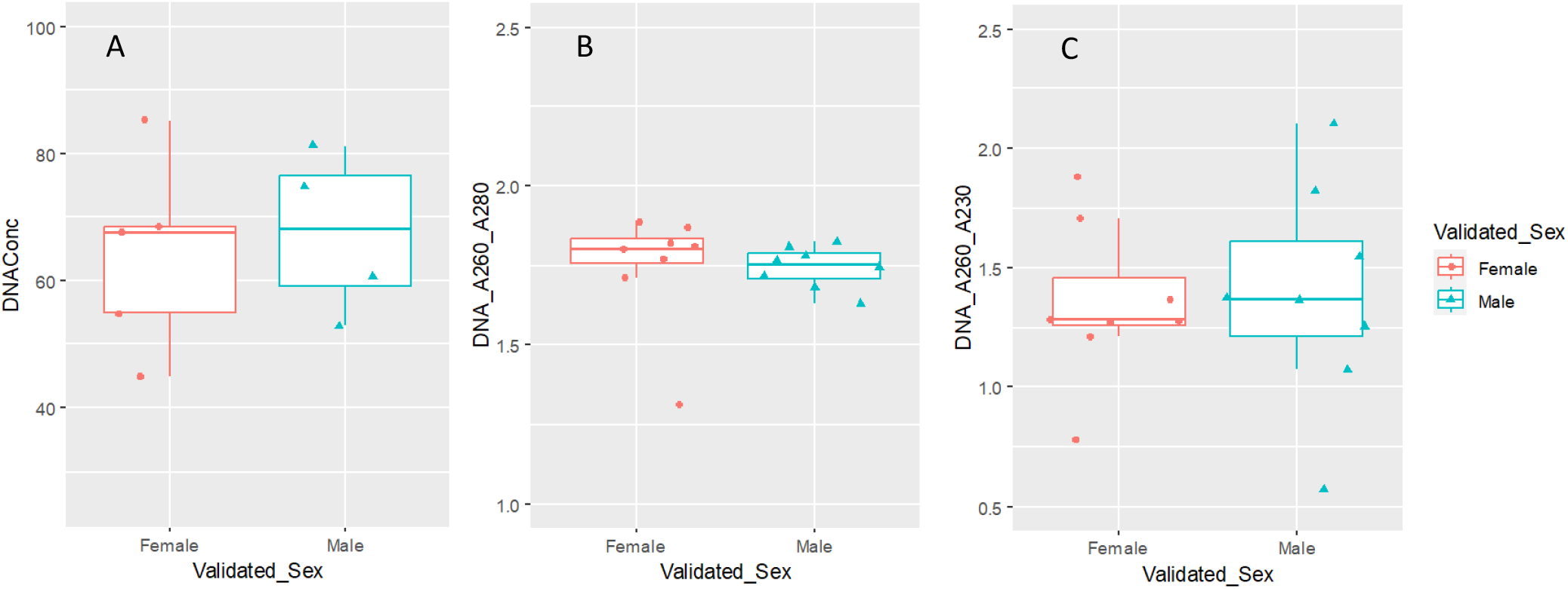
Boxplot jitter plots showing the effect of sex on DNA concentration (A), A260/A280 DNA ratio (B), and A260/A230 DNA ratio (C) in shortnose sturgeon red blood cell samples stored at -80 °C for 8 years (n = 8 for each sex). All three t-tests: df=14, p>0.80.

**Figure 4.**
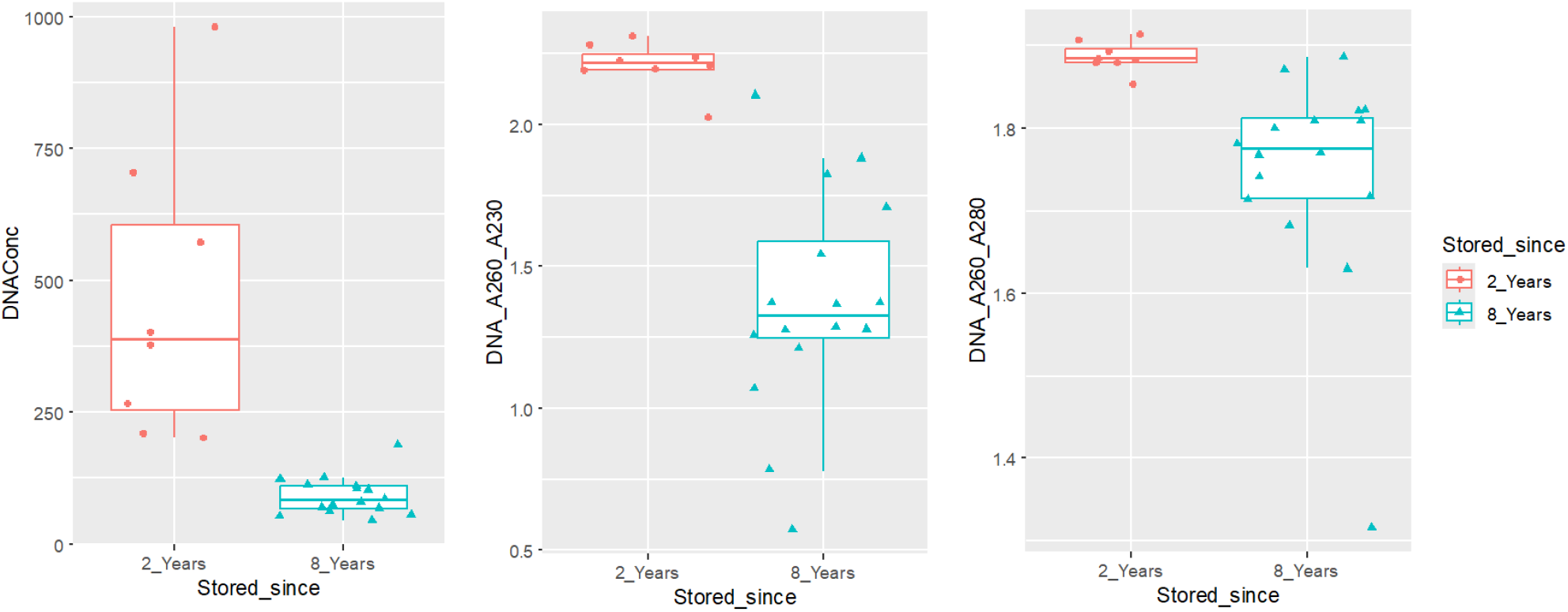
Boxplot jitter plots showing the effect of storage duration on DNA concentration (A), A260/A280 DNA ratio (B), and A260/A230 DNA ratio (C) in shortnose sturgeon red blood cell samples stored at -80 °C for 2 or 8 years (n = 16 for each duration). All three t-tests: df=22, p<<0.001.

**Figure 5.**
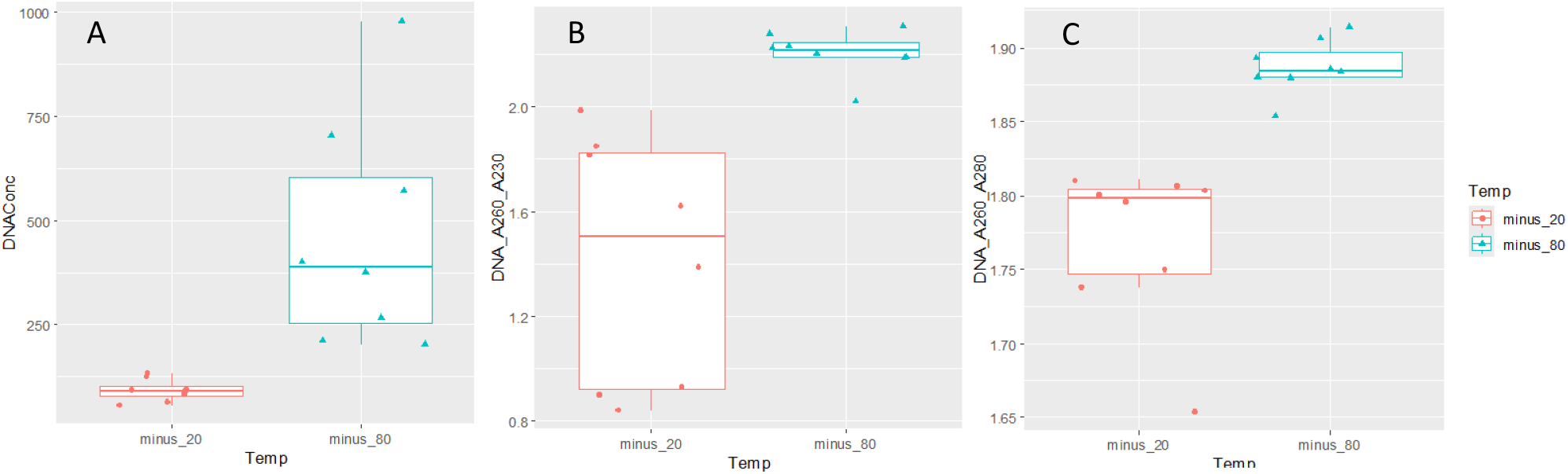
Boxplot jitter plots showing the effect of storage temperature on DNA concentration (A), A260/A280 DNA ratio (B), and A260/A230 DNA ratio (C) in male shortnose sturgeon red blood cell samples stored for 2 years at -20 °C or -80°C (n = 8 for each temperature). All three t-tests: df=14, p<0.002.

## 4. Discussion

This is the first test of Multiplex PCR used to discriminate between male and female sturgeon. Multiplex PCR allows multiple assays to be used at the same time that to save costs and samples. We incorporated an internal control in our Multiplex PCR approach as a safeguard against misinterpretation of males by lack of bands appearing at SSM4 for and ALLWSex2 loci. The purpose of developing and evaluating an internal control (IC) is to monitor the entire detection process, from amplification to final results. Internal controls (ICs) typically involve the use of a separate PCR assay included in the same tube as the target assay [34,35]. An internal control amplifies a nucleic acid target sequence that is invariably present, such as an endogenous housekeeping gene or an exogenous target sequence added with the PCR reagents [36].

The IC serves to monitor the efficiency of each reaction, ensuring that both amplification and detection are functioning effectively, with minimal PCR inhibition that could adversely affect the final results [34,35,37]. If the internal positive control is detected while the target sequence is not, this indicates that the amplification reaction was successful, suggesting that the target sequence is either absent or present at a copy number too low to be detected. Several factors can lead to false negative results, including errors in sample extraction or thermocycler malfunction. However, the most common cause of assay failure is PCR inhibition. By implementing a robust internal control, researchers can enhance the reliability of their PCR assays and ensure accurate detection of target sequences.

Our results clearly shows that RBCs provide an excellent platform to sex shortnose sturgeon using DNA markers. They are superior to fin-clips because they are less likely to be contaminated during sampling and digest faster for PCR analyses. We have also validated SSM4 as another sex-specific marker for shortnose sturgeon. The presence of the SSM4 marker is not surprising, as shortnose sturgeon has been identified phylogenetically as a member of the Acipenserid Atlantic clade based on both microsatellite DNA [38] and mitochondrial genomes [39]. Use of the mtDNA marker provides a check on sample integrity as the absence of both sex markers, SSM4 and ALLWSex2, are used to identify males. In our study mtDNA band occurred in all of our samples confirming sample integrity.

The fish sampled by Litvak’s lab in 2016 were not earmarked for genetic analysis as no markers had been developed until 2021.The RBC samples were duplicates for a stable isotope study on this species. Clearly, storage temperature had an impact on DNA degradation. While flash freezing followed by low temperature storage (−80°C) is the gold standard for long-term storage of tissues for genetic analysis [40,41] samples stored at -20C yielded accurate delineation of sex. This suggests a high degree of sensitivity with this approach and the potential to use it with samples frozen for longer periods. This is particularly important for fieldwork where we do not often have immediate access to liquid nitrogen Dewars or -80°C freezers.

We tested the Multiplex PCR approach with these samples to see if it was robust in sexing fish samples stored for long periods at different temperatures and to look at DNA degradation in long-term storage. As expected, we did see an effect of temperature and storage length. However, the A260/A280 and A260/A230 DNA ratios were high enough to interpret the DNA gels. Validation of our approach on these samples now provides confidence to determine sex for archival samples from this collection and others that have not been validated with other sexing techniques. This will provide important data on sex ratios, spawning biomass and sex differences in distribution of shortnose sturgeon. This is very important to confirm the trajectory of shortnose sturgeon populations over time. This advance will also provide sturgeon farmers an opportunity to detect sex of their fish early in the production cycle.

## Author Contributions

All authors were involved in all stages of completing the research, working on the results and writing the manuscript.

## Institutional Review Board Statement

MTA animal care permit to 103699 to MKL

## Collection permit

MKL—DFO licence 330697.

## Acknowledgments

We would like to thank the DFO Mactaqauc for caring for the shortnose sturgeon used in this study and Alex Giroux for work in MKL’s lab.

